# Genome-scale modeling reveals regulation of human metabolism by the histone deacetylase SIRT1

**DOI:** 10.1101/2025.09.03.673976

**Authors:** Jordi Roma Pi, Jean-Marc Alberto, Justine Paoli, Okan Baspinar, Rosa-Maria Guéant-Rodriguez, Jean-Louis Guéant, Almut Heinken

**Affiliations:** UMRS Inserm 1256 nGERE (Nutrition-Genetics-Environmental Risks), Institute of Medical Research (Pôle BMS)- University of Lorraine, Vandœuvre-les-Nancy, France; University Regional Hospital Nancy, FHU ARRIMAGE, Department of Biochemistry and Molecular Biology, National Reference Laboratory for Inherited Metabolic Diseases; Joan Klein Jacobs Center for Precision Nutrition and Health, Cornell University, Cornell University, Ithaca, USA

**Keywords:** Genome-scale modeling, metabolic regulation, microbiome, SIRT1/butyrate

## Abstract

Genome-scale metabolic models are powerful tools for predicting metabolic fluxes, yet regulatory mechanisms are typically outside their scope. Here, we present a genome-scale modeling framework that integrates transcriptional regulation by the histone deacetylase SIRT1 into human metabolism. By combining a curated regulatory network with the Recon3D metabolic reconstruction, we developed a continuous modeling framework that simulates graded regulatory influences on metabolic fluxes. The model captures known metabolic effects of SIRT1, including enhanced fatty acid oxidation and gluconeogenesis and suppressed glycolysis, across various tissues and dietary conditions. Through cell culture experiments, we quantified the dose-dependent inhibition of SIRT1 by butyrate, a microbiome-derived metabolite. After, incorporating this relationship into the model and found good agreement between experimental metabolomics measurements and in silico predictions. This is the first model to integrate a histone deacetylase and its inhibitor into a genome-scale metabolic framework, enabling simulation of host–microbiome regulatory crosstalk. Our approach provides a dynamic, systems-level tool to explore the regulation of human metabolism and offers insights into how diet and microbial activity influence host metabolic states.

## Introduction

Human metabolism is tightly regulated through signaling and regulatory pathways that control the balance between catabolic and anabolic pathways and respond to the availability of nutrients such as carbohydrates and fatty acids (Chandrasekaran & Price, 2010; Gut & Verdin, 2013; Zhu & Thompson, 2019). Inherited metabolic disorders produce disruptions in metabolic regulation that encompass more metabolic pathways than those directly linked to the mutated gene (Guéant *et al*, 2022; Heinken *et al*, 2024; Wiedemann *et al*, 2024). The disruption of metabolic homeostasis play also a prominent role in the development of noncommunicable complex diseases including obesity, metabolic syndrome, and type 2 diabetes (Diamanti *et al*, 2022; Morgan *et al*, 2016; Nagarajan *et al*, 2022). The underlying mechanisms involve the altered expression of enzymes through genomic and epigenomic mechanisms. An illustrative example is the genomic and epigenomic changes produced by the decreased expression of the histone deacetylase SIRT1 observed in inherited disorders of cobalamin (vitamin B12) metabolism (IDCM) (Gueant *et al*, 2022). Of note, such a decreased expression of SIRT1 is also a key molecular hallmark of obesity and overnutrition. The metabolic fluxomic wide consequences of the altered expression of SIRT1 in IDCM and complex metabolic diseases are not known, given the experimental limitations of the currently available fluxomic techniques.

Mechanisms through which metabolism is controlled include transcriptional regulation, signaling, epigenetics, and posttranslational modifications (Chung *et al*, 2021). Epigenetics refers to inheritable biochemical changes that affect gene expression without modifying the DNA sequence (Gut & Verdin, 2013). One important target of the epigenetic machinery is the packaging of histones and DNA into chromatin, which controls the accessibility of the DNA sequence (Gut & Verdin, 2013). For instance, acetylation of histones, thought to increase the accessibility of DNA, is carried out by histone acetyltransferases (HATs) and reversed by histone deacetylases (HDACs) (Gut & Verdin, 2013). NAD-dependent HDACs, deemed sirtuins, respond to changes in nutritional and energy status by sensing NAD status (Gut & Verdin, 2013). Sirtuin 1 (SIRT1) fulfills an important role in regulating energy metabolism and its expression is induced by factors such as caloric restriction or fasting (Kosgei *et al*, 2020). Besides deacetylation of histones, it also targets nonhistone proteins including peroxisome proliferator-activated receptor- coactivator-1α (PGC1α), Fork Head Box O1 transcription factor (Foxo1), and Peroxisome proliferator activated receptor α (PPARα). Through targeting these transcription factors, activation of SIRT1 induces fatty acid oxidation and gluconeogenesis, and suppresses glycolysis (Kosgei *et al*., 2020). In adipocytes, SIRT1 promotes insulin sensitivity, lipid metabolism, mitochondrial biogenesis, and white to brown adipocyte differentiation (Kosgei *et al*., 2020; Majeed *et al*, 2021). Taken together, expression of SIRT1 is protective against obesity and metabolic syndrome (Kosgei *et al*., 2020). Consequently, obese individuals have been shown to have reduced SIRT1 expression, and SIRT1 levels increase after weight loss (Mengozzi *et al*, 2022; Rappou *et al*, 2016).

An important factor in regulation of human metabolism mediated by diet is the gut microbiome (Sonnenburg & Backhed, 2016). Accordingly, changes in composition and function of the gut microbiome have been implicated in complex metabolic diseases related to metabolic syndrome including obesity, type 2 diabetes, and nonalcoholic fatty liver disease (Delzenne *et al*, 2020; Takeuchi *et al*, 2023; Tilg *et al*, 2021). Besides other metabolic functions, gut microbes the short-chain fatty acids acetate, propionate, and butyrate regulate host metabolism through multiple routes (Sonnenburg & Backhed, 2016). For instance, butyrate inhibits SIRT1 in a dose-dependent manner (Pant *et al*, 2019), hence linking gut microbiome metabolism to host metabolic regulation.

Investigating the complex mechanisms linking human metabolism and regulatory pathways requires a systems-level approach. Systems biology methods aim to comprehensively characterize the components of a biological system, such as a cell or organism, through computational methods such as network modeling. Constraint-Based Reconstruction and Analysis (COBRA) is a systems biology approach relying on detailed, manually curated genome-scale reconstructions of metabolism (O’Brien *et al*, 2015). Genome-scale reconstructions can be converted into mathematical models through the implementation of constraints such as mass-charge balance and the nutrient environment (O’Brien *et al*., 2015). Through the most basic constraint-based modeling method, flux balance analysis (FBA), a space of feasible fluxes is computed under the basic assumption that metabolic networks are in a steady state where the concentration of metabolites remains constant over time (Orth *et al*, 2010). Condition-specific metabolic models can be generated by integrating context-specific omics data such as transcriptomics (Sen & Oresic, 2023). The COBRA approach has also been applied to the prediction of metabolic interactions between the host and the gut microbiome (Heinken *et al*, 2021). For instance, personalized gut microbiome models can also be integrated with a whole-body model of human metabolism, enabling the prediction of the effects of microbial metabolites on distant organs such as the brain (Thiele *et al*, 2020).

Despite the many advantages of COBRA, one drawback is that in their basic form, most genome-scale reconstructions only account for metabolic pathways and lack regulatory mechanisms such as signaling, transcriptional regulation, posttranslational modification, or epigenetics (Chung *et al*., 2021). This limitation is significant as it restricts the model’s ability to predict cellular responses to external stimuli such as signaling molecules, which play a key role in fine-tuning metabolic fluxes. For instance, while the integrated whole-body model of human metabolism and the gut microbiome successfully recapitulated that microbial butyrate serves as the carbon source for the colonocyte (Thiele *et al*., 2020), it did not capture the regulatory effects of butyrate, e.g., its inhibition of host histone deacetylases. Moreover, while thousands of genome-scale reconstructions of metabolism have been built for all domains of life (Heinken *et al*, 2023; Ye *et al*, 2022), genome-scale reconstructions of regulation are less common. One notable exception is a recent genome-scale reconstruction of the transcriptional regulatory network in placenta (Paquette *et al*, 2024).

To overcome this limitation, modeling frameworks have been developed to integrate regulatory mechanisms with metabolic reconstructions (Chung *et al*., 2021; Cruz *et al*, 2020). In the first such work, transcriptional regulatory networks (TRNs) were integrated with genome-scale metabolic reconstructions by formulating regulatory rules as discrete Boolean logic where gene regulatory states are represented as binary on/off conditions, and integrating them with the gene-protein-reaction associations in the metabolic reconstruction (Covert *et al*, 2004). Dedicated implementations of FBA deemed regulatory flux balance analysis (rFBA) (Covert *et al*., 2004) and steady-state regulatory flux balance analysis (SR-FBA) (Shlomi *et al*, 2007) enabled predicting the effect of regulatory rules on metabolic fluxes. While useful for simulating the presence or absence of gene activity, this binary approach simplifies the complex nature of gene regulation, which can vary continuously and dynamically in response to environmental stimuli. To overcome this issue, Probabilistic Regulation of Metabolism (PROM) instead implemented continuous regulatory rules that enforced metabolic fluxes to be proportional to the probability of their encoding genes (Chandrasekaran & Price, 2010). Notably, while several works have integrated transcriptional regulatory networks with metabolic reconstructions, representations of epigenetic mechanisms in genome-scale models are less common (Chung *et al*., 2021). A recent developed tool, MEWpy, further facilitated integrating regulatory and metabolic data within genome-scale metabolic models (Pereira *et al*, 2021). Recently, genome-scale metabolic modeling has been proposed as a tool for modeling the interactions between HDACs, their inhibitors, and metabolism as it accounts for gene-protein-reaction relationships as well as nutrient availability and metabolite levels (i.e., of HDAC inhibitors) (King *et al*, 2021).

In this study, we present a regulatory genome-scale modeling framework that integrates a transcriptional regulatory network of the histone deacetylase SIRT1, deemed iSIRT1, with the human genome-scale metabolic reconstruction Recon3D. By linking SIRT1 to metabolism, we provide a comprehensive framework to explore its multifaceted role in metabolic regulation. A key feature of iSIRT1 is the incorporation of the inhibition of SIRT by butyrate, a dietary metabolite with known effects on metabolic health, into the model. This addition allows us to model indirect effects of microbial metabolites on human metabolism through regulatory pathways with SIRT1 as a case study.

## Methods

### Construction of iSIRT1

A systematic computational approach was used to construct iSIRT1, based on interaction data retrieved via the pypath Python library (https://pypath.omnipathdb.org). This framework interfaces with OmniPath, a literature-curated resource of signaling and regulatory interactions (Türei D), and enabled automated extraction, filtering, and preprocessing of relevant interactions.

### Interaction Retrieval

Interactions involving SIRT1 were queried from OmniPath, a comprehensive resource that integrates manually curated databases including SIGNOR2. The query included protein-protein interactions (PPIs), transcriptional regulations, and metabolic associations. The query identified seven transcription factors (Supplementary Table S1) as direct targets of SIRT1, from which downstream regulatory interactions were retrieved. After filtering for high-confidence, experimentally supported entries, a curated interaction set of 494 transcription factor-gene regulatory interactions was retained for downstream analysis (Supplementary Table S1).

### Mapping Pathways and Regulatory Layers

To extend the regulatory network, a depth-first search algorithm was used to identify all downstream metabolic genes regulated by the seven transcription factors (TF). These genes were then mapped to metabolic pathways using Recon3D annotations. The resulting hierarchical network linked SIRT1 to key metabolic subsystems, including fatty acid oxidation, gluconeogenesis, and glycolysis. The final curated network, deemed iSIRT1, included 487 transcriptionally regulated metabolic genes and 2,296 targeted metabolic reactions in Recon3D (Brunk *et al*, 2018) (Supplementary Table S2).

### Retrieval of Regulatory Rules

Following the construction of the hierarchical network linking SIRT1, transcription factors, and metabolic genes, regulatory rules were formulated to capture the influences exerted by TF on their target genes. Each metabolic gene was systematically analyzed to determine whether its regulation was positive (activation) or negative (repression).

### Rule Formulation

For each metabolic gene, the combined effect of all influencing TF was defined using logical operators. Initially, Boolean logic was applied with the standard connectors "AND," "OR," and "NOT" to describe the interactions. However, Boolean logic assumes binary states (0 or 1) for gene expression, which oversimplifies the nuanced dynamics of regulatory networks. For example, when a gene is influenced by an activating TF (expression = 1.0) and a repressing TF (expression = 0.0), the traditional Boolean "AND" operation calculates the minimum (min(1.0, 0.0) = 0.0), inaccurately implying complete repression.

To address these limitations, a mean-based computation strategy was implemented for the "AND" and "OR" operators. Under this approach, the output reflects the average influence of all relevant TF. For the example, the mean is calculated as (1.0 + 0.0) / 2 = 0.5, representing a neutral regulatory influence. This adjustment provides a more realistic depiction of regulatory dynamics, particularly when conflicting influences are present.

### Implementation in MEWpy

To integrate transcriptional regulation into constraint-based modeling, we used the MEWpy toolbox (Pereira *et al*., 2021) and extended its SR-FBA framework to support continuous-valued regulatory logic. This modification allows gene expression to influence metabolic fluxes in a graded rather than binary manner. Details of the implementation and representative logic rules are provided in Supplementary Notes Section 1 and Supplementary Table S1.

### Integration of iSIRT1 with Metabolic Reconstruction

Following the formulation of regulatory rules, iSIRT1 was integrated with the Recon3D genome-scale metabolic model (Brunk et al., 2018), obtained from the Virtual Metabolic Human Database (Noronha A, 2019). Genes not represented in Recon3D were excluded to ensure compatibility. The resulting integrated model, iSIRT1_HumanMet, 8 regulators, 487 transcriptionally regulated metabolic genes, and 2,296 targeted metabolic reactions (within a total of 10,607 reactions in Recon3D).

Technical implementation details, including modified files and symbolic evaluation strategies, are described in Supplementary Methods, Section 2.

### Simulations

Flux Balance Analysis (FBA) (Orth *et al*., 2010) and Flux Variability Analysis (FVA) (Mahadevan & Schilling, 2003) were performed on the continuous regulatory-metabolic model iSIRT1_Recon3D to evaluate flux distributions under varying regulatory and environmental conditions. All simulations were executed in steady state using the modified MEWpy framework with the CPLEX solver (version 22.x) and COBRApy (Ebrahim *et al*, 2013) as the modeling backend.

### Robustness analysis

To determine the metabolic model’s flexibility under differing regulatory constraints, SIRT1 expression levels were systematically varied from 0.0 to 1.0 in increments of 0.1. At each expression level, FBA was performed using biomass production as the sole objective function. Fluxes were computed across all reactions, and subsystem-level flux summaries were generated to evaluate transcriptional regulation effects on metabolism. FVA was used to determine the minimum and maximum allowable fluxes through each reaction at each SIRT1 level.

Dietary environments were incorporated by constraining the uptake rates of nutrient exchange reactions. Five distinct dietary regimes (high-protein, high-carbohydrate, high-fat, ketogenic, and Western diet) were implemented by modifying previously defined in silico diet constraints (Heinken *et al*, 2013). Uptake rates used to simulate each diet are provided in Supplementary Table S3.

### Construction of tissue-specific models

Tissue-specific genome-scale metabolic models were generated using the FASTCORE algorithm (Pacheco & Sauter, 2018; Vlassis *et al*, 2014), based on transcriptomic data from the GTEx Project (Consortium *et al*, 2020). To define the set of core reactions for each tissue, gene expression values were first binarized and mapped to the gene-protein-reaction (GPR) associations in the global iSIRT1 model. Reactions linked to expressed genes were then used to construct each tissue-specific model with FASTCORE, ensuring both metabolic functionality and biological relevance.

Each resulting context-specific model was further extended with a SIRT1-centered regulatory network. This enabled us to perform steady-state continuous regulatory flux balance analysis (srFBA) simulations across a range of SIRT1 expression levels (from 0.0 to 1.0 in increments of 0.1). All simulations used the *biomass_reaction* as the objective function, with standard uptake constraints unless otherwise noted.

This modeling framework builds on the approach established by Pacheco & Sauter (2018), who integrated GTEx transcriptomic data with genome-scale metabolic models to investigate human tissue metabolism. The complete reconstruction pipeline, including scripts to derive tissue-specific models from Recon3D and apply regulatory constraints, is available at https://github.com/almut-heinken/ngereSysBio/tree/main/iSIRT1_regulatoryModeling.

### Cell culture experiments

To explore the effects of butyrate on SIRT1 expression and cellular metabolism, in vitro experiments were performed using human Caco-2 intestinal epithelial cells. Cells were cultured from a frozen batch (passage 7) and initially seeded into T25 flasks for 7 days, followed by 7 days of expansion in petri dishes. For treatment, cells were divided into 30 petri dishes (10 per condition group) and incubated for 72 hours with increasing concentrations of sodium butyrate (0, 0.3, 1.0, 3.0, and 9.0 mM).

After incubation, supernatants were collected and frozen at –80 °C for LDH release and free DNA quantification, while cell pellets were harvested for downstream analyses. SIRT1 protein expression was measured by Western blot and normalized to TBP expression. In parallel, metabolomics analyses were performed to quantify intracellular concentrations of butyrate and TCA cycle intermediates via LC-MS. RNA and DNA were extracted for gene expression and quantification assays, and enzymatic assays were conducted to evaluate SIRT1 activity.

A full description of the experimental setup, including treatment groups, sample processing, and measured endpoints, is available in Supplementary Methods Section 1 and illustrated in Supplementary Figure S1.

### Parameterization of iSIRT1_HumanMet with butyrate inhibition

The data from the cell culture experiments were used to derive a quantitative regression model linking butyrate flux to SIRT1 expression. To facilitate computational modeling, we converted the experimental butyrate uptake into flux units (mmol/gDW/h), considering treatment volume, duration, cell number, and dry weight assumptions (Supplementary Methods, Section 1). The regression model, which showed a statistically significant inverse linear relationship (R² = 0.510, p = 0.004; Supplementary Figure S2), was then used to estimate SIRT1 expression from flux values in subsequent simulations.

### Validation with Caco-2 Metabolomics Data

To assess the agreement between model predictions and experimental data, we used intracellular metabolite concentrations measured from Caco-2 cells treated with varying concentrations of butyrate (0, 0.3, 1, 3, and 9 mM). A total of 15 metabolites had been measured, of which 2-Hydroxyglutarate and 2-Ketoglutarate were excluded due to absence in the metabolic reconstruction, resulting in 13 metabolites used for validation.

Building on top of the existing Recon3D reactions, we introduced custom demand reactions for each of the 13 metabolites of interest. For each butyrate condition, the extracellular concentration (in mM) was translated into an estimated uptake flux (mmol/gDW/h) and applied as a model constraint.

We then performed SIRT1-regulated Flux Balance Analysis (srFBA) for each metabolite and condition, optimizing for the maximum possible flux through its corresponding demand reaction. Each simulation accounted for the respective SIRT1 expression level, enabling regulatory influence to be incorporated directly into the optimization.

Simulation results were compared to all corresponding experimental replicates at the matching butyrate concentration. Statistical comparison between predicted and observed values was performed using the Mann–Whitney U test and Spearman correlation. The full set of comparisons is presented in Supplementary Figure S3.

### Personalized modeling of host inhibition by butyrate

To incorporate microbial butyrate production into host metabolic simulations, we retrieved butyrate fluxes predicted by microbiome community models (Shaaban *et al*, 2024) that had been built for a cohort of infant and maternal gut microbiomes from the AGORA2 resource (Heinken *et al*., 2023) through the mgPipe (Heinken & Thiele, 2022) workflow. To enable integration of predicted microbiome fluxes into the host genome-scale reconstruction Recon3D, the standard unit in mgPipe of mmol/person/day (Heinken & Thiele, 2022) was converted to the unit convention used in Recon3D of mmol/gDW/h. Microbial biomass was estimated for each infant and adult based on estimated body weight, with a scaling factor applied to estimate microbial mass in grams dry weight (gDW). The microbial butyrate flux was then adjusted using a factor accounting for the wet-to-dry weight ratio of human tissue. These adjusted fluxes were used to estimate butyrate production, which was further integrated into downstream models, including the regression model derived from the Caco-2 cell experiments (Supplementary Methods, Section 1). This methodology enabled us to model the effects of personalized microbial butyrate production on host SIRT1 expression, providing a foundation for subsequent simulations of host metabolism. For further details on the flux conversion process and assumptions, see Supplementary Methods, Section 3.

## Results

We present iSIRT1_HumanMet, a predictive model that integrates a SIRT1-centered transcriptional regulatory network into the Recon3D genome-scale metabolic framework. To simulate graded regulatory influences, we implemented a continuous logic framework, allowing real-valued gene activity states to constrain metabolic fluxes within the COBRA environment. This enabled a dynamic exploration of how incremental changes in SIRT1 expression shape the metabolic landscape. Furthermore, by parameterizing the model with experimental data from Caco-2 cells, we incorporated butyrate-induced inhibition of SIRT1, enabling simulation of microbiome-derived regulatory effects. We demonstrate below that iSIRT1_HumanMet highlights key pathway-level flux changes as a function of SIRT1 activity, with emphasis on central carbon metabolism, mitochondrial function, and lipid utilization.

### Development of an Integrated Regulatory-Metabolic Model Workflow

We aimed to construct a unified modeling framework capable of simulating the graded regulatory effects of SIRT1 on human metabolism at genome scale. To achieve this, we developed a continuous regulatory-metabolic model by integrating a curated transcriptional network of SIRT1, deemed iSIRT1_HumanMet (in silico SIRT1) and accounting for 8 regulators, with the Recon3D metabolic reconstruction (Methods). The resulting hybrid model, which couples regulatory and metabolic layers through continuous transcriptional logic, was deemed iSIRT1_HumanMet and included 487 regulated genes linked to 2,296 targeted metabolic reactions (out of 10,607 reactions in Recon3D) and enabled condition-specific simulations of SIRT1-mediated metabolic shifts. Logical regulatory rules were formulated to represent the influence of SIRT1 on transcription factors and metabolic genes and were incorporated into the iSIRT1 model using a continuous logic formulation. This approach allowed gene expression levels to act as dynamic flux constraints, in contrast to existing Boolean-based methods such as PROM (Chandrasekaran & Price, 2010).

## Figure Legends

summarizes the overall modeling workflow, from interaction retrieval to rule formulation and integration with the metabolic network. Taken together, iSIRT1 successfully captures continuous regulatory influences and provides a robust platform to explore how transcriptional regulation modulates metabolic behavior across a range of biological conditions.

### Features of the Integrated Regulatory-Metabolic Model

We next characterized the structure and coverage of the integrated model, iSIRT1_HumanMet, which combines transcriptional regulation and genome-scale metabolism. The base metabolic framework, Recon3D, includes 10,607 reactions, 5,839 metabolites, and 1,886 genes. Within this network, we incorporated a SIRT1-centered regulatory layer comprising eight transcriptional regulators, with SIRT1 as the upstream node.

The downstream regulatory cascade includes FOXO1, FOXO3, PPAR*α*, PPAR*γ*, PPARGC1*α*, TP53, and EP300, all of which are known to be directly modulated by SIRT1 (Ding *et al*, 2017; Feige *et al*, 2008; Gerhart-Hines *et al*, 2007; Kosgei *et al*., 2020; Rodgers & Puigserver, 2007). These regulators form a hierarchical network of 494 curated interactions, ultimately controlling the expression of 487 metabolic genes. The regulated genes influence 2,296 targeted metabolic reactions within the Recon3D network, highlighting the extensive regulatory reach of SIRT1 within the metabolic network.

In summary, iSIRT1_HumanMet enables simulation of regulatory control across nearly half of the metabolic reactions in Recon3D, providing a comprehensive framework for studying SIRT1-mediated metabolic adaptation.

### SIRT1 Activity Rewires Central Metabolic Fluxes

To evaluate the regulatory influence of SIRT1 on metabolism, we applied Flux Variability Analysis (FVA) across a range of SIRT1 expression levels (0.0 to 1.0, in 0.1 increments) using iSIRT1_HumanMet. This allowed systematic quantification of how transcriptional activation of SIRT1 alters the feasible flux space of all reactions in Recon3D.

SIRT1 activation resulted in increased flux capacity through enzymes involved in gluconeogenesis and fatty acid β-oxidation, consistent with a shift toward endogenous glucose production and enhanced lipid catabolism (Figure 2A). In contrast, several glycolytic enzymes exhibited reduced flux, suggesting decreased reliance on glucose as a primary energy source under high SIRT1 activity. These model predictions are consistent with experimental findings showing that SIRT1 promotes oxidative metabolism through PGC-1α activation and represses glycolysis under metabolic stress or fasting conditions (Canto & Auwerx, 2012; Gerhart-Hines *et al*., 2007; Rodgers & Puigserver, 2007).

**Figure 1.**
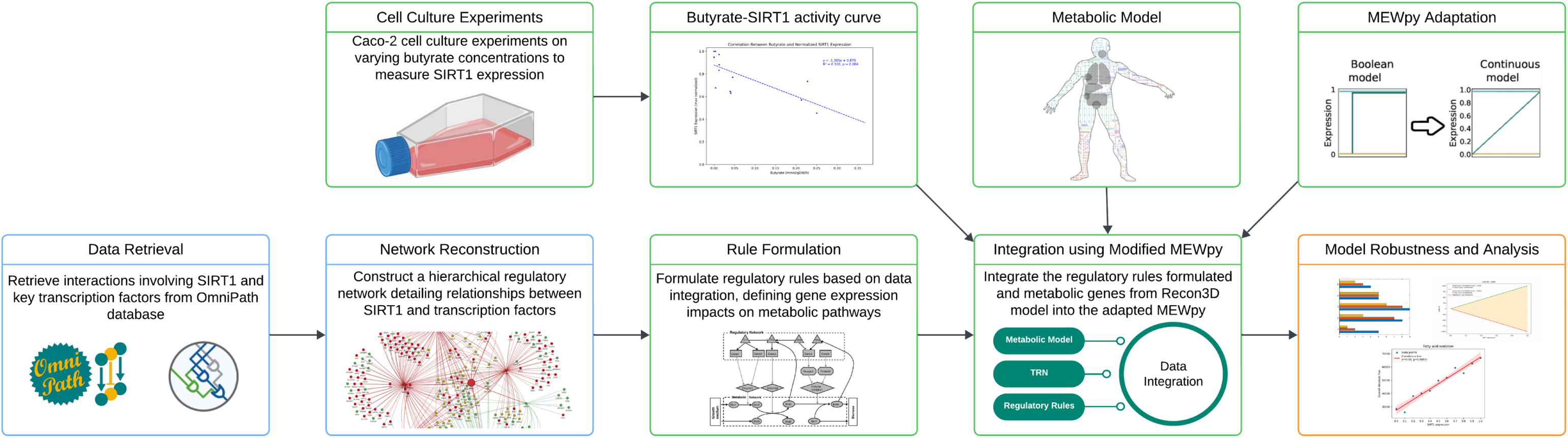
Workflow for integrating continuous regulatory logic with a genome-scale metabolic model. Illustration of the pipeline used to construct the iSIRT1_HumanMet model, integrating transcriptional regulation with metabolism. The workflow combines cell culture experiments, regulatory network reconstruction, and continuous regulatory logic implemented in a modified MEWpy framework. Interactions involving SIRT1 were retrieved from OmniPath, followed by formulation of gene-specific rules and integration into the Recon3D metabolic model. The final model was evaluated through robustness analysis to assess regulatory impact on metabolic behavior. Abbreviations: VMH, Virtual Metabolic Human; TF, transcription factor; srFBA, steady-state regulatory Flux Balance Analysis.

**Figure 2.**
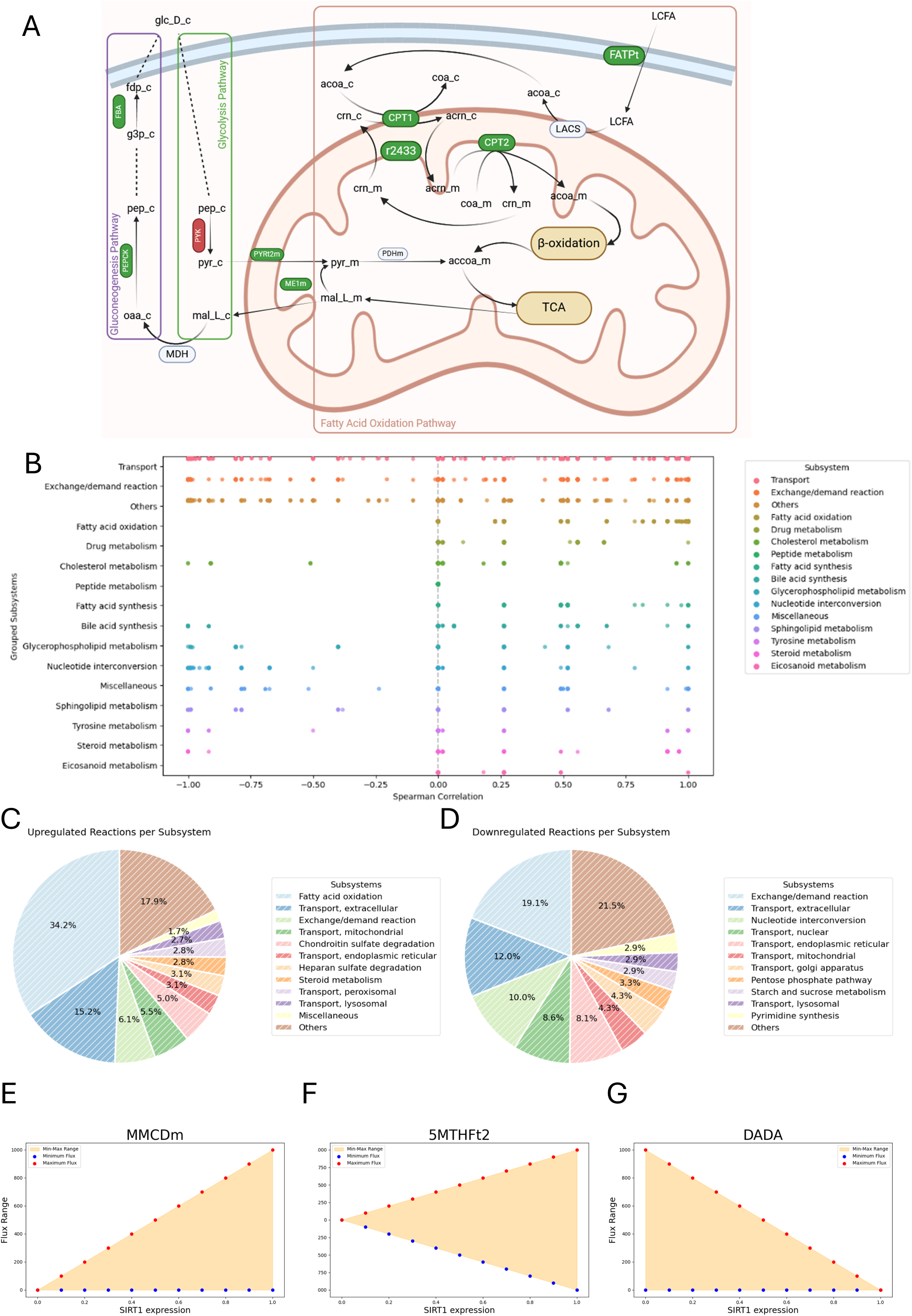
Predicted effects of SIRT1 activity on metabolic reactions. Regulatory impact of SIRT1 expression on cellular metabolism predicted by iSIRT1_HumanMet, as inferred from flux balance simulations using the integrated regulatory-metabolic model. Both global and subsystem-specific trends are visualized to highlight pathways influenced by varying SIRT1 levels. A. Schematic representation of selected metabolic pathways influenced by SIRT1 activity. Predicted regulatory effects on key enzymes are color-coded (green: upregulated, red: downregulated), covering glycolysis, the TCA cycle, and fatty acid oxidation, with compartmental transport steps shown. Metabolite abbreviations follow Virtual Metabolic Human (VMH) naming conventions. B. Spearman correlation coefficients between SIRT1 expression and predicted reaction fluxes across the model, grouped and color-coded by metabolic subsystem. Each dot represents a reaction, with its correlation strength indicating the direction and magnitude of SIRT1’s effect. C. Distribution of upregulated reactions across metabolic subsystems. Fatty acid oxidation, transport reactions, and extracellular exchanges represent the largest contributions. D. Distribution of downregulated reactions across metabolic subsystems. Exchange reactions and nucleotide interconversion are among the most affected. E. FVA result for MMCDm (Methylmalonyl Coenzyme A Decarboxylase, Mitochondrial), a key reaction in folate/1-carbon metabolism. The allowable flux range expands with increasing SIRT1 expression, indicating enhanced flexibility in this pathway. F. FVA result for 5MTHFt2 (5-Methyltetrahydrofolate Transport via Anion Exchange), another component of folate/1-carbon metabolism. The reaction shows increasing flux range with higher SIRT1 expression, suggesting potential activation or derepression. G. FVA result for DADA (Deoxyadenosine Deaminase), part of the nucleotide interconversion subsystem. The flux range decreases as SIRT1 expression rises, indicating repression of this pathway under high SIRT1 activity.

Global correlation analysis revealed strong positive associations between SIRT1 expression and fluxes through fatty acid oxidation, transport, and exchange/demand reactions (Figure 2B). In contrast, fluxes in glycolysis and nucleotide interconversion reactions were negatively correlated with SIRT1 activity, reinforcing a global regulatory shift toward energy-efficient lipid metabolism.

Subsystem-level analysis showed that 34.2% of upregulated reactions belonged to the fatty acid oxidation pathway, followed by extracellular transport (15.2%) and exchange/demand (13.7%) (Figure 2C). Downregulated reactions primarily affected exchange/demand (21.5%), extracellular transport (19.1%), and nucleotide interconversion (12.0%) (Figure 2D). These patterns are consistent with SIRT1’s established role in promoting mitochondrial energy metabolism and suppressing anabolic or rapid turnover pathways.

Targeted analysis of individual reactions revealed increased flux through carnitine palmitoyltransferases (CPT1 and CPT2), which are essential for mitochondrial fatty acid uptake and oxidation. Similarly, phosphoenolpyruvate carboxykinase (PEPCK), a key gluconeogenic enzyme, exhibited elevated flux at higher SIRT1 expression levels (Figure 2A). These findings support previous reports that SIRT1 enhances mitochondrial lipid oxidation via PGC-1α and upregulates gluconeogenesis via deacetylation of FOXO1 and PGC-1α in various experimental conditions (Canto & Auwerx, 2012; Gerhart-Hines *et al*., 2007; Rodgers & Puigserver, 2007), including in myocardium of rat pups from mothers subjected to a deficient diet in folate and vitamin B12 during pregnancy and lactation (Garcia *et al*, 2011).

To further explore mechanistic links between SIRT1 and one-carbon metabolism (1CM), we examined three reactions with notable flux changes across the SIRT1 gradient (Figures 2E–G). Methylmalonyl-CoA decarboxylase (MMCDm) (Figure 2E) displayed an increasing flux range with rising SIRT1 levels. This enzyme catalyzes the decarboxylation of methylmalonyl-CoA to propanoyl-CoA and is a key component of the B12-dependent anaplerotic TCA pathway, particularly relevant for the oxidation of odd-chain fatty acids. The observed increase in flexibility supports a model in which SIRT1 facilitates mitochondrial integration of alternative carbon sources, including B12-dependent substrates.

In Figure 2F, the 5-methyltetrahydrofolate transport via anion exchange (5MTHFt2) reaction showed increased bidirectional flux capacity under high SIRT1 expression. Although the exact transporter modeled may not correspond directly to the reduced folate carrier (RFC/SLC19A1), this reaction serves as a proxy for mitochondrial folate exchange, a critical element of 1CM. Enhanced flexibility in this transport step suggests that SIRT1 promotes redistribution of folate intermediates, potentially supporting redox homeostasis, methylation potential, or nucleotide biosynthesis under stress.

Finally, Figure 2G illustrates a decrease in the flux range of D-amino acid dehydrogenase (DADA) with increasing SIRT1, indicating a repression of amino acid catabolism in favor of more energy-efficient substrates such as lipids. This aligns with SIRT1’s broader role in resource conservation and metabolic streamlining under low-nutrient conditions.

In conclusion, simulations using iSIRT1 reveal a coordinated reprogramming of metabolism upon SIRT1 activation, favoring fatty acid oxidation and gluconeogenesis while repressing glycolysis. These results align with experimental data and support the use of iSIRT1 as a predictive tool for dissecting SIRT1-mediated metabolic regulation.

### Dietary influence on metabolic regulation mediated by SIRT1

To assess how dietary inputs influence SIRT1-regulated metabolism, we performed Flux Balance Analysis (FBA) on iSIRT1_HumanMet under five dietary conditions, Western, High-Carbohydrate, High-Fat, Ketogenic, and High-Protein (Methods, Supplementary Table S3). FBA simulations were performed across SIRT1 expression levels ranging from 0.0 to 1.0 in 0.2 increments.

SIRT1 activation consistently increased flux through fatty acid oxidation pathways, with the strongest responses observed under High-Fat and Ketogenic diets (Figure 3A). This trend aligns with experimental studies reporting that SIRT1 promotes mitochondrial β-oxidation and lipid utilization under energy-demanding conditions through the deacetylation of PGC-1α and activation of fatty acid oxidation genes (Canto & Auwerx, 2012; Rodgers & Puigserver, 2007). Conversely, pathways associated with nucleotide interconversion and glycine, serine, and threonine metabolism showed decreasing flux with higher SIRT1 expression, especially under Western and High-Carbohydrate diets. This suggests a reduction in energetically costly anabolic pathways as SIRT1 promotes metabolic efficiency (Kosgei *et al*., 2020).

**Figure 3.**
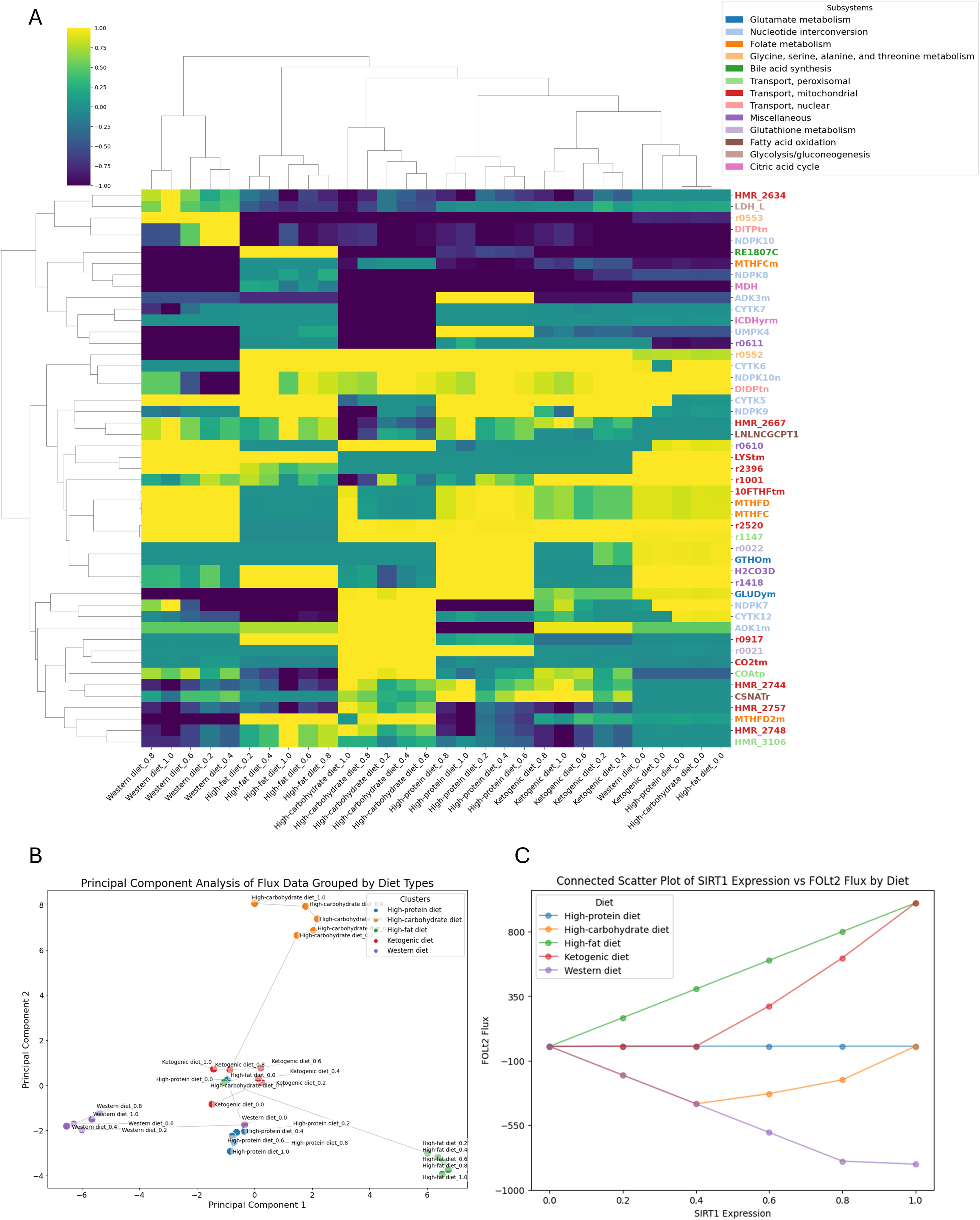
Reaction fluxes and PCA across diet conditions and SIRT1 expression levels. Metabolic profiles computed in Recon3D on varying SIRT1 expression on five different dietary regimes. Reactions were simulated across five diet types and varying SIRT1 levels, and their flux responses were analyzed globally and individually. A. Heatmap of normalized reaction fluxes across combinations of five diet types and SIRT1 expression levels (0.0 to 1.0). Each row represents a metabolic reaction, color-coded by subsystem. Columns correspond to specific diet-expression combinations. Hierarchical clustering highlights patterns of co-regulated reactions across conditions. B. Principal Component Analysis (PCA) of predicted flux distributions, grouped by diet type. Each point represents a flux profile at a specific SIRT1 expression level, and clustering indicates that diet composition influences the principal flux variation more strongly than SIRT1 level alone. C. Connected scatter plot showing the predicted flux of FOL2 (dihydrofolate reductase) as a function of SIRT1 expression under each diet type. Results demonstrate that dietary context modulates the sensitivity of folate metabolism to SIRT1 activity.

To evaluate global metabolic trends, we applied Principal Component Analysis (PCA) to the simulated flux profiles (Figure 3B). At low SIRT1 expression (0.0), metabolic differences between diets were minimal, as indicated by overlapping clusters. As SIRT1 expression increased, diets became more separated in PCA space, indicating that SIRT1 amplifies diet-specific metabolic responses. Although High-Fat and Ketogenic diets did not fully cluster together, they showed similar directions of separation along Principal Components 1 and 2, reflecting overlapping adaptations despite contrasting macronutrient composition. Interestingly, High-Carbohydrate also followed a similar trend, suggesting partial convergence in metabolic strategy.

In conclusion, simulations with iSIRT1_HumanMet demonstrated that diet composition modulates the extent and direction of SIRT1-dependent metabolic reprogramming. Lipid-rich diets further enhance fatty acid catabolism in response to SIRT1, while carbohydrate-based diets exhibit stronger repression of nucleotide and amino acid pathways. These results reinforce the role of SIRT1 as a metabolic integrator of nutritional state and suggest mechanistic links between diet, transcriptional regulation, and metabolic flexibility (Canto & Auwerx, 2012; Garcia *et al*., 2011; Kosgei *et al*., 2020; Rodgers & Puigserver, 2007).

### Tissue-specific regulation of metabolism by SIRT1

SIRT1 regulates key metabolic pathways in a tissue-specific manner, with its effects varying across organs depending on physiological context and transcriptional activity (Gerhart-Hines *et al*., 2007; Li *et al*, 2011). To explore these differences, tissue-specific metabolic models were generated using transcriptomic data (see Methods).

Simulations were run across SIRT1 expression levels from 0.0 to 1.0, revealing considerable tissue-to-tissue variability in active metabolic components (Table 2). Principal Component Analysis of the resulting flux profiles showed distinct clustering of functionally related tissues, indicating a strong regulatory influence of SIRT1 on metabolic specialization (Figure 4C). FVA further highlighted SIRT1-mediated differences in glycolysis, oxidative phosphorylation, and nucleotide metabolism across tissues (Figure 4D). SIRT1 knockout simulations led to widespread flux disruptions, with pronounced effects on oxidative phosphorylation, glycolysis/gluconeogenesis, and fatty acid oxidation (Supplementary Figure S4), reinforcing its role in systemic metabolic regulation.

**Figure 4.**
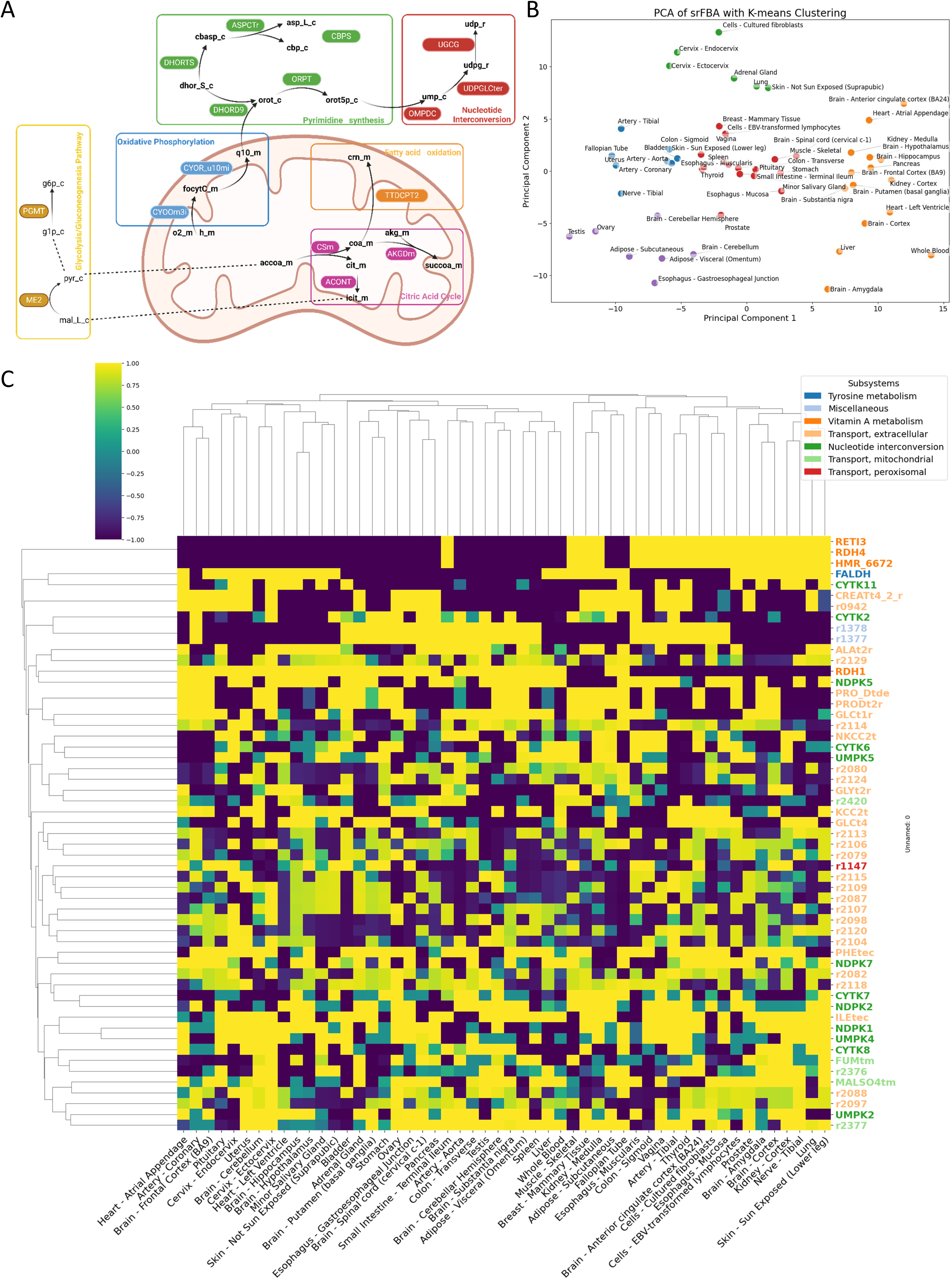
Tissue-specific metabolic models and flux variability under SIRT1 regulation. Generation and analysis of tissue-specific regulatory-metabolic models incorporating SIRT1 activity. Shown are the effects of SIRT1-mediated transcriptional regulation on metabolic flexibility across human tissues. A. Table summarizing the number of reactions, metabolites, genes, and target genes across the reconstructed tissue-specific models, including counts of shared and unique components. These models were derived using transcriptomic data integrated into a Recon3D-based framework. B. Schematic overview of selected metabolic reactions predicted to be regulated by SIRT1, grouped by subsystem (e.g., oxidative phosphorylation, nucleotide interconversion, pyrimidine synthesis, and fatty acid oxidation). The diagram highlights cross-compartment fluxes affected by regulatory activity. C. Principal Component Analysis (PCA) of predicted flux distributions from steady-state regulatory FBA (srFBA) across 54 tissues. Each point represents a tissue-specific flux profile, with colors indicating k-means clustering. The plot reveals functional clustering of tissues based on their metabolic response to SIRT1 regulation. D. Heatmap showing the flux variability of selected reactions across all tissues. Each row corresponds to a reaction (color-coded by subsystem), and each column represents a tissue. Color intensity reflects the magnitude of flux variability, revealing tissue-specific regulatory influence on core metabolic pathways.

**Table 1.** Examples of Boolean regulatory rules used in the metabolic-regulatory model. Each rule defines the transcriptional regulation of a metabolic gene based on upstream transcription factors. Logical operators include AND, OR, and NOT. The full set of curated rules is provided in Supplementary Table S2.

| Gene Name | Regulatory Rule |
| --- | --- |
| G1 | TF1 |
| G2 | TF1 OR TF2 |
| G3 | TF3 AND NOT TF4 |

**Table 2.**
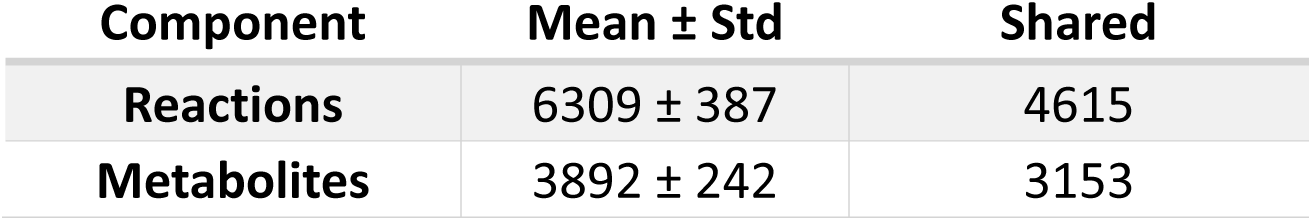
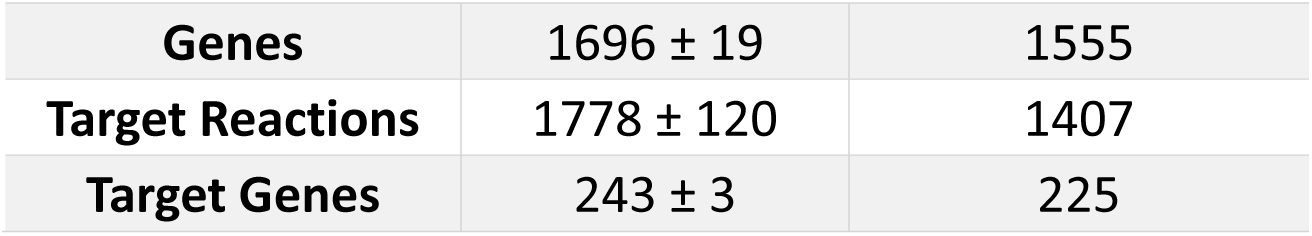
Summary of Core and Variable Components Across Tissue-Specific Metabolic Models. Summary statistics of reactions, metabolites, genes, and regulatory targets across the tissue-specific metabolic models reconstructed using GTEx transcriptomic data and the Recon3D framework. The table reports the mean ± standard deviation of the number of elements per tissue model, as well as the number of components shared across all tissues.

| Component | Mean ± Std | Shared |
| --- | --- | --- |
| Reactions | 6309 ± 387 | 4615 |
| Metabolites | 3892 ± 242 | 3153 |
| **Genes** | 1696 ± 19 | 1555 |
| **Target Reactions** | 1778 ± 120 | 1407 |
| **Target Genes** | 243 ± 3 | 225 |

| Component | Mean ± Std | Shared |
| --- | --- | --- |
| Reactions | 6309 ± 387 | 4615 |
| Metabolites | 3892 ± 242 | 3153 |
| Genes | 1696 ± 19 | 1555 |
| Target Reactions | 1778 ± 120 | 1407 |
| Target Genes | 243 ± 3 | 225 |

Taken together, these tissue-specific simulations demonstrate that SIRT1 contributes to metabolic flexibility and energy homeostasis by modulating central metabolic pathways in a tissue-dependent manner. An experimental illustration if the key role of SIRT1 in the dysregulation of metabolic homeostasis in myocardium, gut and brain, but not in liver, in rat pups from mothers subjected to a deficient diet in folate and vitamin B12 during pregnancy and lactation and IDCM fibroblasts (Garcia *et al*., 2011; Ghemrawi *et al*, 2013; Gueant *et al*., 2022; Melhem *et al*, 2016; Pooya *et al*, 2012; Pourié *et al*, 2015).

### Model predictions of butyrate-mediated inhibition of SIRT1 agree with independent metabolomics data

Butyrate inhibits SIRT1 activity in a dose-dependent manner (Pant *et al*., 2019). The inhibitory effect of butyrate on SIRT1 expression was experimentally determined in Caco-2 intestinal epithelial cells across a gradient of physiologically relevant concentrations (0–9 mM) (Methods). Western blot analysis confirmed a concentration-dependent reduction in SIRT1 protein levels, which was modeled using a linear regression to relate butyrate flux to SIRT1 expression. This relationship was incorporated into the model to simulate regulatory flux responses under different butyrate exposures using steady-state regulatory FBA (Methods).

To evaluate the model’s biological validity, we compared predicted secretion fluxes against experimental metabolomics data collected from the same Caco-2 cultures that had not been used during the generation of the in-silico model. Out of 15 quantified intracellular metabolites, 13 could be mapped to exchange reactions in the model; 2-hydroxyglutarate and 2-ketoglutarate were excluded due to their absence from the model structure.

For each butyrate condition, the model produced a single simulated flux value per metabolite, which was directly compared to the three corresponding experimental replicates (without transformation). Most measured metabolites displayed strong agreement between simulations and experimental results (Supplementary Figure S3). Statistically significant Spearman correlations (p < 0.05) were found for 11 of 13 metabolites, including lactate (ρ = 0.88), succinate (ρ = 0.94), and 3-hydroxypropionate (ρ = 0.84), supporting the model’s ability to reproduce key metabolic trends under increasing butyrate levels.

While the regulatory mechanism in the model was implemented via changes in SIRT1 expression, the observed flux differences likely reflect a combination of both butyrate-driven effects and SIRT1-dependent regulation. Additional targeted experiments or simulations would be needed to disentangle these two components.

### Simulation of SIRT1 Regulation by Gut Microbial Butyrate Fluxes

To evaluate the potential regulatory effects of microbiome-derived butyrate on host SIRT1 expression, we leveraged predicted butyrate secretion fluxes that had been predicted in silico for the gut microbiomes of a cohort of infants during the first year of life (Shaaban *et al*., 2024). These predictions had been generated from microbiome community models built through the AGORA2 resource (Heinken *et al*., 2023) and using shotgun metagenomic profiles of 20 infants (sampled at 5 days, 1 month, 6 months, and 1 year) and 13 mothers.

Flux values, originally reported in units of mmol/person/day, were converted to mmol/gDW/h using microbial biomass estimates and age-specific average host body weights (see Supplementary Table S4 and Supplementary Methods, Section 4). This normalization enabled integration into the same modeling framework used for the Caco-2 experiments.

We applied the linear regression derived from the in vitro butyrate–SIRT1 experiment (Supplementary Figure S2) to translate predicted butyrate fluxes into inferred SIRT1 expression levels for each individual sample (Methods). As shown in Figure 5A, infant microbiomes exhibited lower butyrate fluxes at early time points, corresponding to higher SIRT1 levels, while increased fluxes at later stages suggested progressive downregulation of SIRT1. Principal Component Analysis of microbial fluxes (Figure 5B) revealed clear age-related clustering, reflecting temporal maturation of microbial metabolism. Differences were particularly driven by time-dependent shifts in short-chain fatty acid metabolism, nucleotide interconversion, and amino acid biosynthesis pathways, suggesting that microbial functional development may influence host regulatory processes such as SIRT1 activity. Taken together, these results support a mechanistic link between microbial butyrate production and host SIRT1 expression during early development and provide insight into how gut microbiome activity may influence host epigenetic and metabolic programming.

**Figure 5.**
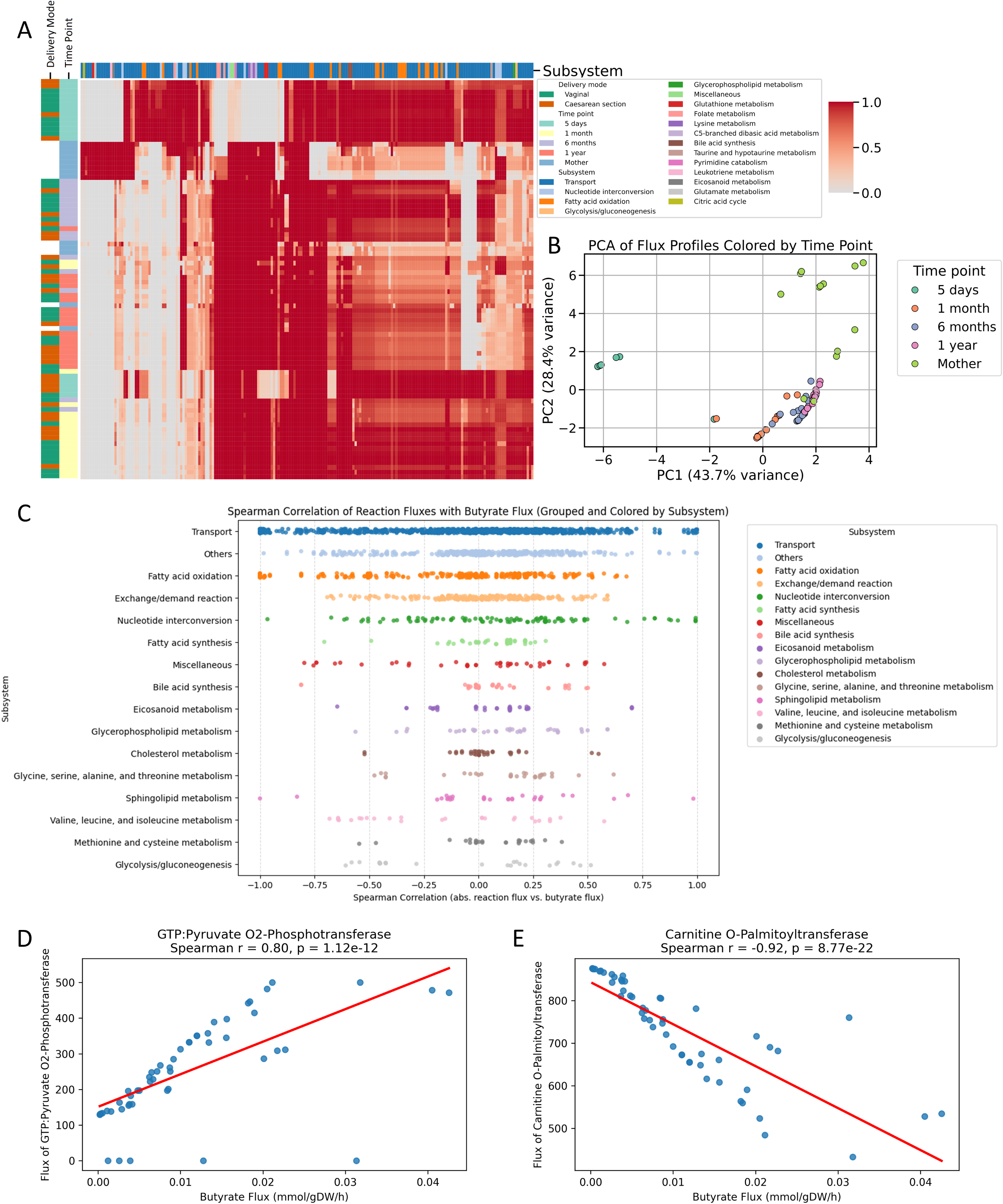
Modulation of human metabolic fluxes by the gut microbial metabolite butyrate via its inhibition of SIRT1 activity. Butyrate production predicted for the gut microbiomes in a cohort of infant and mothers (Shaaban et al., 2024) had been used to contextualize iSIRT1_HumanMet. Shown are the predicted human fluxes as a function of SIRT1 inhibition via sample-specific butyrate uptake flux. A. Heatmap of normalized human reaction fluxes across samples from 20 infants (sampled at 5 days, 1 month, 6 months, and 1 year) and 13 mothers. Rows represent individual microbial reactions grouped by metabolic subsystem (color-coded), and columns correspond to individual samples. Side bars indicate delivery mode (vaginal or cesarean) and time point. B. Principal Component Analysis (PCA) of the microbial flux profiles. Each point represents one sample and is colored by time point. The separation along PC1 reflects temporal differences in microbial function, distinguishing early infancy from later stages and maternal profiles. C. Spearman correlation between absolute butyrate flux and individual microbial reaction fluxes across all samples. Reactions are grouped and color-coded by metabolic subsystem. Positive and negative correlations highlight functional pathways most associated with microbial butyrate production. D. Scatter plot showing a strong positive correlation between butyrate flux and the flux of GTP-pyruvate O₂-phosphotransferase, a reaction involved in energy metabolism (Spearman r = 0.80, p = 1.12e−12). E. Scatter plot showing a strong negative correlation between butyrate flux and the flux of carnitine O-palmitoyltransferase, a key enzyme in fatty acid oxidation (Spearman r = −0.92, p = 8.77e−22). This suggests an inverse relationship between butyrate production and lipid catabolism in microbial communities.

## Discussion

This study presents a novel genome-scale metabolic model that integrates SIRT1’s regulatory effects on human metabolism, providing a more dynamic and biologically accurate representation of regulatory influence on metabolic networks. We combined experimental data, regulatory network reconstruction, and mechanistic modeling to investigate how SIRT1 activity, modulated by microbial butyrate, impacts metabolic fluxes across tissues, diets and developmental stages. By extending the MEWpy framework to support continuous-valued regulation, we captured graded transcriptional responses and identified context-specific shifts in key pathways including gluconeogenesis, fatty acid oxidation, and nucleotide interconversion. This integrative approach offers new insight into the interplay between microbial metabolites, gene regulation and human metabolism.

Unlike previous approaches relying on Boolean logic (Chung *et al*., 2021; Covert *et al*., 2004; Shlomi *et al*., 2007), our model incorporates a continuous regulatory framework, enabling nuanced, graded regulatory effects that better reflect biological processes. This advancement allows for a more precise simulation of metabolic shifts under different physiological conditions. Consistent with experimental findings, the model captures SIRT1’s role in promoting energy-efficient processes such as fatty acid oxidation and gluconeogenesis while suppressing glycolysis (Canto & Auwerx, 2012; Kosgei *et al*., 2020), reinforcing its function in oxidative metabolism during energy-stressed states such as fasting. By integrating continuous regulatory signals into metabolic simulations, this work refines the depiction of cellular metabolism’s dynamic complexity and provides a robust framework for further investigation into regulatory mechanisms and metabolic disease modeling.

Our integrated model of SIRT1 regulation and metabolism successfully reproduces known SIRT1-regulated pathways, notably the promotion of fatty acid oxidation and gluconeogenesis, both of which enhance metabolic efficiency under energy stress (Canto & Auwerx, 2012; Kosgei *et al*., 2020). Activation of SIRT1 enhances mitochondrial function, promotes lipid utilization, and suppresses glycolysis, thereby shifting metabolic preference toward oxidative metabolism (Gerhart-Hines *et al*., 2007; Gueant *et al*., 2022; Rodgers & Puigserver, 2007). These regulatory effects are particularly relevant in the context of metabolic syndrome and type 2 diabetes, where disrupted energy metabolism contributes to insulin resistance, ectopic lipid accumulation, and systemic inflammation (Ding *et al*., 2017; Jalgaonkar *et al*, 2022; Tilg *et al*., 2021). By restoring oxidative metabolic programs, SIRT1 activation has been proposed as a promising therapeutic strategy for improving metabolic health (Barber *et al*, 2022). The ability of our model to capture these fundamental SIRT1-mediated metabolic shifts highlights its potential for simulating metabolic interventions and exploring regulatory mechanisms underlying complex metabolic diseases.

Beyond these primary pathways, the model revealed additional SIRT1-influenced pathways, including nucleotide metabolism and amino acid interconversion, suggesting broader regulatory roles. For example, the observed shift away from nucleotide metabolism under high-carbohydrate conditions may reflect an adaptive response orchestrated by SIRT1 to conserve cellular resources during energy stress. These secondary effects expand the know influence of SIRT1 beyond energy homeostasis, implicating it in the maintenance of biomass and redox balance, processes that are also disrupted in obesity, insulin resistance, and inflammatory conditions (Kosgei *et al*., 2020; Majeed *et al*., 2021). Thus, targeting SIRT1 could have broader therapeutic benefits than previously recognized, affecting not only energy metabolism but also cellular biosynthesis and stress resilience.

Tissue-specific modeling further demonstrated that SIRT1 exerts distinct regulatory effects across different organs. In adipose tissue, SIRT1 promotes lipid metabolism, mitochondrial biogenesis, and insulin sensitivity (Majeed *et al*., 2021; Mengozzi *et al*., 2022), while in the liver, it enhances gluconeogenesis and fatty acid oxidation, supporting systemic glucose and lipid homeostasis (Kosgei *et al*., 2020; Rodgers & Puigserver, 2007). These findings are consistent with literature emphasizing SIRT1’s context-dependent regulatory roles and suggest that therapeutic modulation of SIRT1 should account for tissue-specific metabolic demands. Targeted activation or inhibition of SIRT1 in specific organs could offer more precise strategies to address IDCM and complex metabolic diseases such as obesity, type 2 diabetes, and nonalcoholic fatty liver disease (Barber *et al*., 2022; Gueant *et al*., 2022; Tilg *et al*., 2021).

The gut microbiome is known to regulate host metabolism through nutrient signaling, immune modulation, and epigenetic mechanisms (Sonnenburg & Backhed, 2016; Tilg *et al*., 2021). Our study demonstrates that microbiome-derived metabolites such as butyrate can modulate SIRT1 expression, providing a mechanistic link between diet, microbial metabolism, and host regulatory networks. Experimental validation in Caco-2 cells revealed a dose-dependent inhibitory effect of butyrate on SIRT1 protein levels, and computational integration of microbiome flux models extended these findings by predicting host SIRT1 regulation across early life stages. The observed developmental increase in microbial butyrate flux and corresponding inferred downregulation of SIRT1 highlight the dynamic interplay between microbial activity and host gene regulation. These findings suggest that microbiome composition and metabolic output could influence critical host pathways involved in energy homeostasis, mitochondrial function, and inflammatory responses (Barber *et al*., 2022; Kosgei *et al*., 2020). Given that dysbiosis and altered short-chain fatty acid profiles have been implicated in metabolic syndrome, obesity, and type 2 diabetes (Delzenne *et al*., 2020; Tilg *et al*., 2021), microbiome-mediated modulation of SIRT1 may represent an important factor in disease susceptibility. Future work integrating longitudinal microbiome, dietary, and host epigenetic data could further elucidate how early-life microbial exposures shape metabolic health trajectories.

Traditionally COBRA models have predominantly relied on binary on/off states to simulate gene regulation (Chung *et al*., 2021; Covert *et al*., 2004; Shlomi *et al*., 2007), which, although effective in some contexts, oversimplify the dynamic nature of gene regulation. In contrast, our continuous framework captures graded regulatory influences, enhancing the physiological realism and predictive capacity of metabolic simulations. This successful integration of SIRT1 regulatory pathways demonstrates the feasibility of combining regulatory and metabolic networks at a higher resolution, paving the way for more sophisticated models that incorporate complex regulation, such as transcriptional, signaling, and epigenetic influences (Chung *et al*., 2021). Although a significant step forward, further work should aim to incorporate additional regulatory layers, including post-translational modifications and stochastic regulatory effects, to more fully realize the potential of regulatory-metabolic reconstructions.

While several works have integrated transcriptional regulatory networks with metabolic reconstructions, representations of epigenetic mechanisms in genome-scale models are less common (Chung *et al*., 2021). In a first work, context-specific models of macrophages were generated from transcriptomics data, and histone H3K27 acetylation data was used to identify genes with high regulatory load (Pacheco *et al*, 2015). Another noteworthy effort is the work by Shen et al., who studied the metabolic dependencies of deacetylase inhibitors by representing cellular histone acetylation through a bulk acetylase and deacetylase reaction (Shen *et al*, 2019). In another recent study, DNA methylation and demethylation reactions were included into the human genome-scale reconstruction Human1, and cancer cell line-specific subnetworks were contextualized with experimentally measured cell line-specific methylation levels (Barata *et al*, 2024). To our knowledge, this is the first study to integrate the regulatory network of a histone deacetylase and its inhibitors into a metabolic reconstruction. Future developments could see the integration of bulk acetylation and deacetylation into iSIRT1_HumanMet.

We acknowledge that iSIRT_HumanMet currently has several limitations. First, the regulatory framework focuses specifically on SIRT1-mediated deacetylation and does not yet incorporate other post-translational modifications or signaling pathways that also influence metabolic regulation (Chung *et al*., 2021). Expanding the model to capture a broader range of microbiome-mediated regulatory interactions, e.g., metabolic targets by other members of the sirtuin family of histone deacetylases (Wu *et al*, 2022), activation of G-protein-coupled receptors (Sonnenburg & Backhed, 2016), or activation of aryl hydrocarbon receptors (Modoux *et al*, 2022) could enhance its predictive capabilities. Including further regulatory targets in human metabolism of microbiome-derived metabolites, e.g., short-chain fatty acids (Delzenne *et al*., 2020; Gut & Verdin, 2013; Sonnenburg & Backhed, 2016), would enhance the model’s ability to simulate the complex microbiome-host metabolic crosstalk.

Second, the reliance on transcriptomics data to infer regulatory activity may not fully capture the dynamic and context-dependent nature of regulatory responses, such as transient signaling events or post-transcriptional modifications (Chung *et al*., 2021) that occur independently of gene expression changes. Incorporating epigenomic data, including histone modifications and DNA methylation profiles (Barata *et al*., 2024; Gut & Verdin, 2013), could further refine the regulatory framework, enabling dynamic simulations of host metabolic responses to epigenetic changes.

Finally, although the use of a continuous regulatory framework represents an improvement over traditional binary Boolean logic (Chandrasekaran & Price, 2010; Covert *et al*., 2004), further refinements are needed to incorporate probabilistic or dynamic regulatory models that better reflect stochastic gene expression and variability across physiological conditions. Addressing these limitations could lead to more comprehensive and physiologically realistic simulations of metabolic regulation in future work. Such advancements would support a holistic, systems-level understanding of human metabolism, particularly in the context of metabolic and inflammatory diseases, and could inform the development of personalized therapeutic strategies targeting the microbiome-epigenome-metabolism axis.

## Author Contributions

A.H., J-L.G., and J.R.P. designed research. J.R.P. built the integrated transcriptional regulatory and metabolic model and performed simulations. R-M.G.R. and J-M.A. designed and performed cell culture experiments. J.P. performed Western blot analyses. O.B. performed metabolomics analyses. J.R.P. and A.H. analyzed results. J.R.P. and A.H. drafted the manuscript. J.R.P., A.H., J-L.G., and J-M.A. revised the manuscript. A.H. and J-L.G. supervised the project. All authors reviewed and approved the final version of the manuscript.

## Data and Code Availability

Scripts, input data that enable reproducing the simulations and analyses, and results of performed simulations are available at https://github.com/almut-heinken/ngereSysBio/tree/main/iSIRT1_regulatoryModeling. The fork of MEWpy (https://github.com/BioSystemsUM/MEWpy) containing functions modified in this study can be found at https://github.com/jordirpUL/MEWpy. The regulatory model (iSIRT1.xml, 26 MB) is available at https://github.com/almut-heinken/ngereSysBio/blob/main/iSIRT1_regulatoryModeling/data/models/sirt1_recon3d/iSIRT1.xml.

---

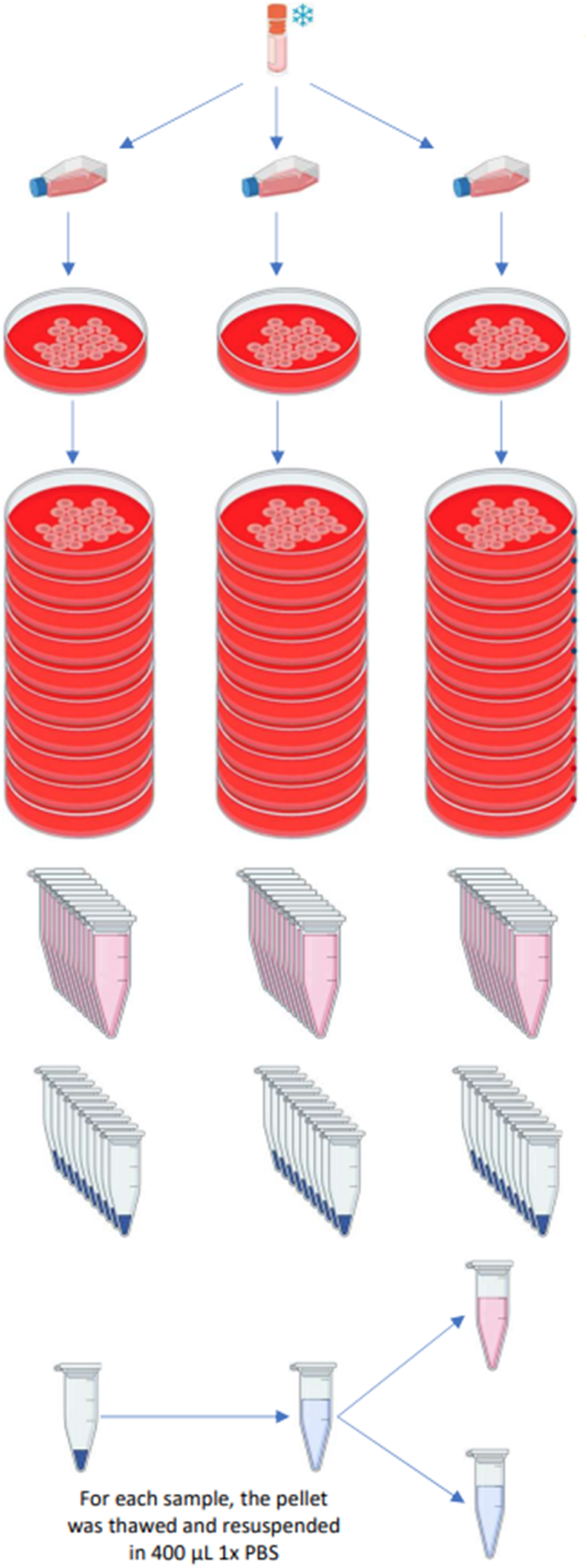

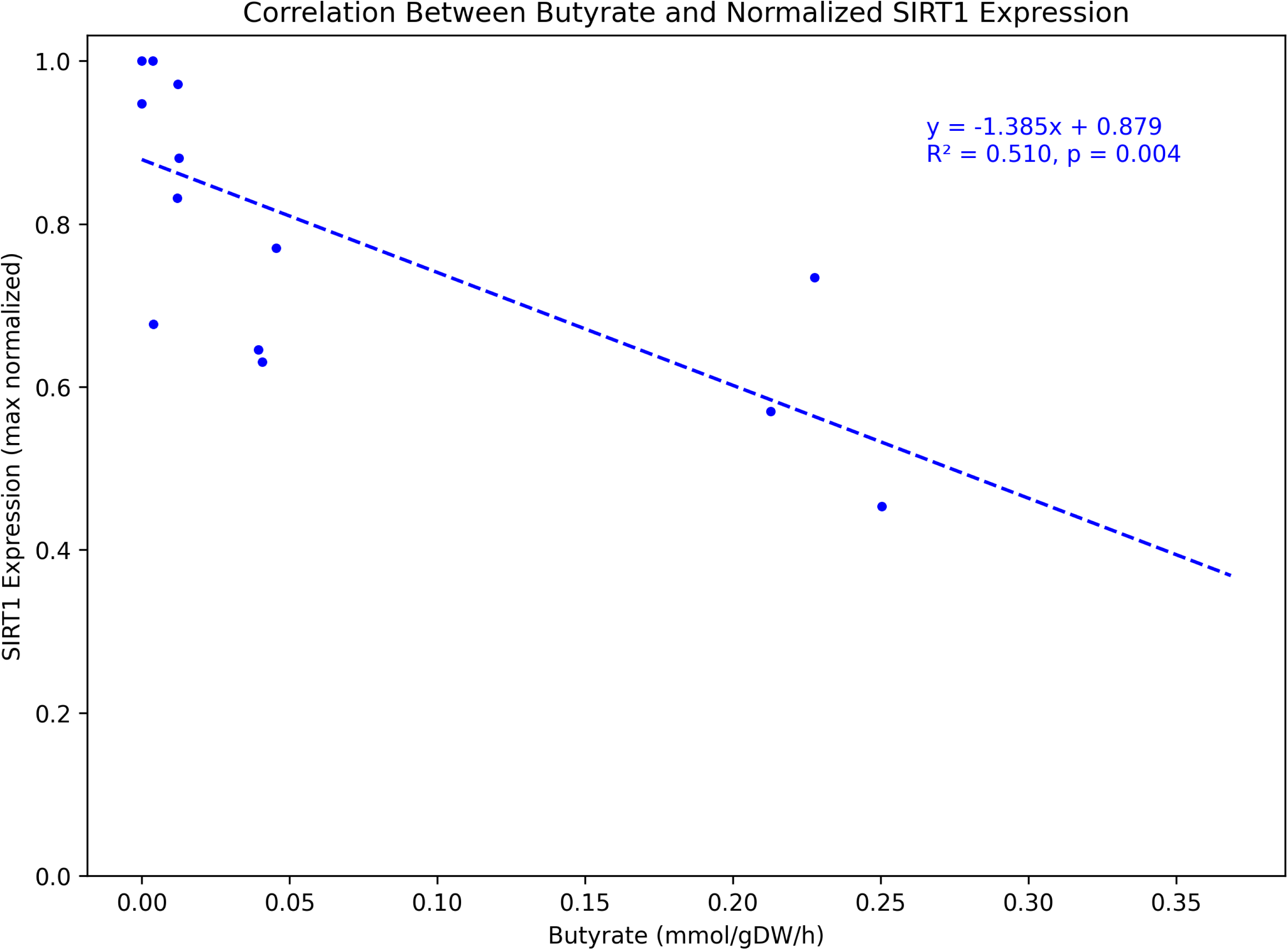

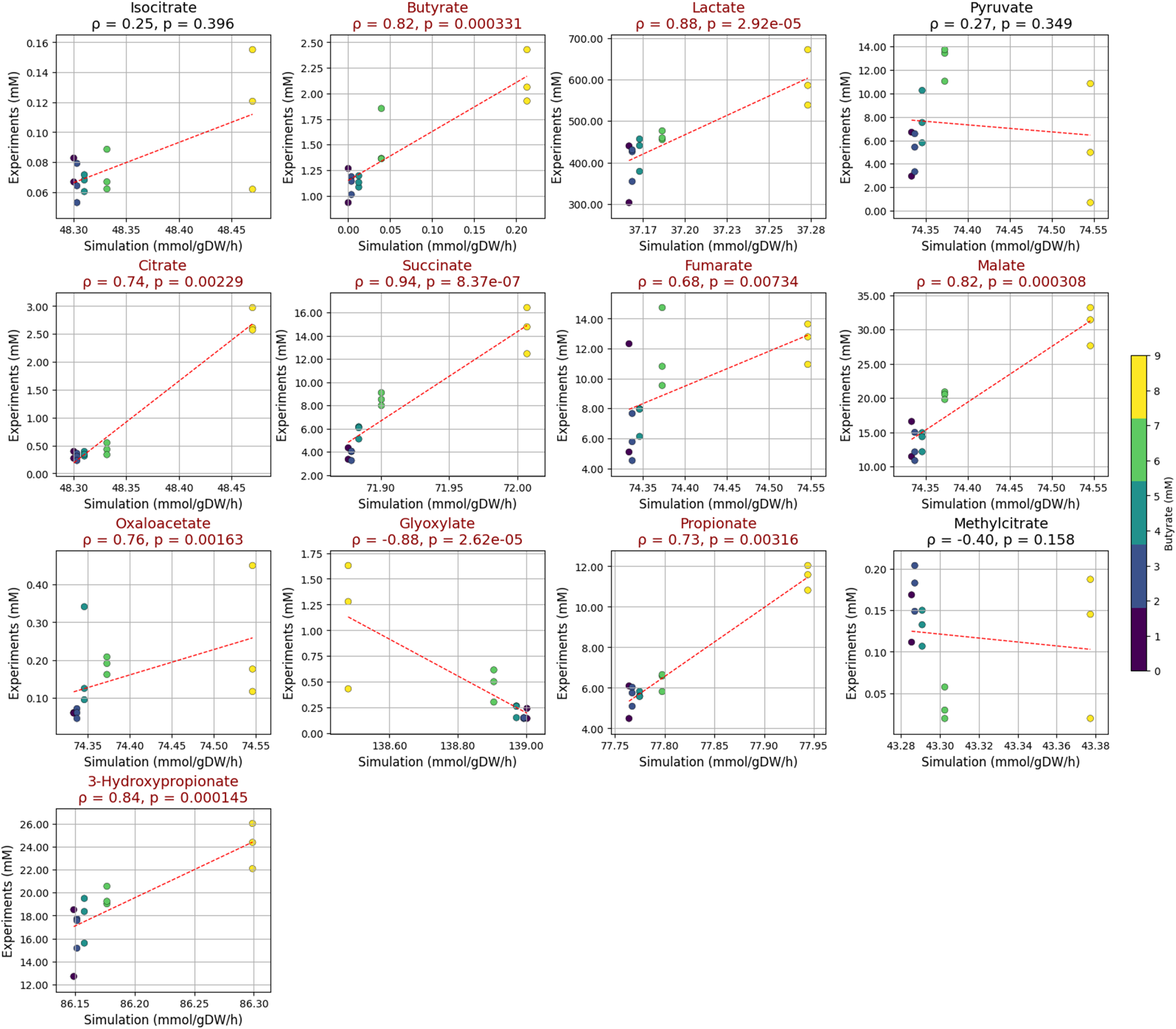

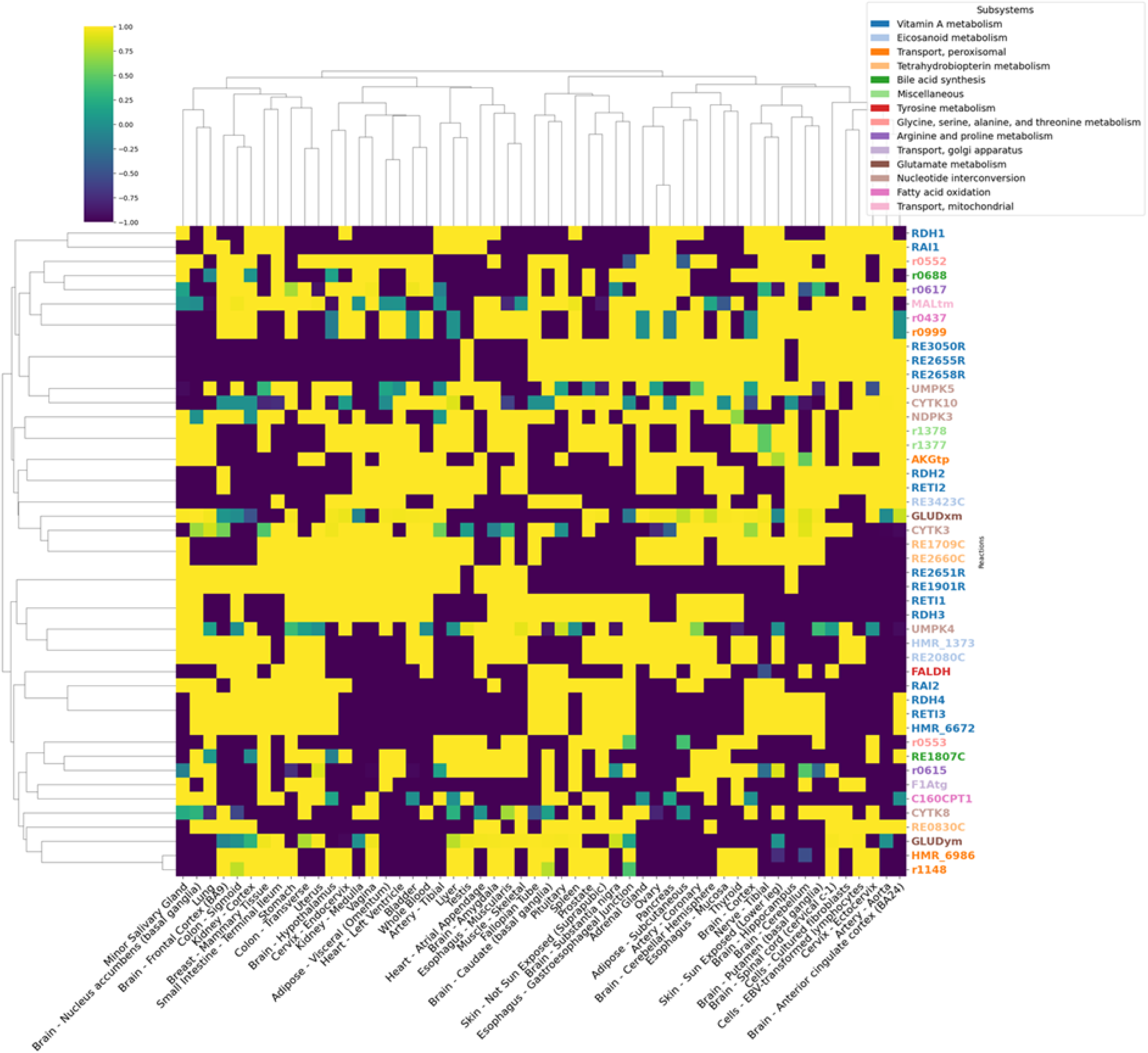

